# Wireless endovascular nerve stimulation with a millimeter-sized magnetoelectric implant

**DOI:** 10.1101/2021.07.06.450036

**Authors:** Joshua C. Chen, Peter Kan, Zhanghao Yu, Fatima Alrashdan, Roberto Garcia, Amanda Singer, C.S. Edwin Lai, Ben Avants, Scott Crosby, Michelle M. Felicella, Ariadna Robledo, Jeffrey D. Hartgerink, Sunil A. Sheth, Kaiyuan Yang, Jacob T. Robinson

**Affiliations:** Department of Bioengineering, Rice University, Houston, TX, USA; Department of Neurosurgery, University of Texas Medical Branch, Galveston, TX, USA; Department of Electrical and Computer Engineering, Rice University, Houston, TX, USA; Applied Physics Program, Rice University, Houston, TX, USA; Neuromonitoring Associates, LLC; Department of Pathology, University of Texas Medical Branch, Galveston, TX, USA; Department of Chemistry, Rice University, Houston, TX, USA; Department of Neurology, UTHealth McGovern Medical School, Houston, TX, USA; Department of Neuroscience, Baylor College of Medicine, Houston, TX, USA

## Abstract

Implanted bioelectronic devices have the potential to treat disorders that are resistant to traditional pharmacological therapies; however, reaching many therapeutic nerve targets requires invasive surgeries and implantation of centimeter-sized devices. Here we show that it is possible to stimulate peripheral nerves from within blood vessels using a millimeter-sized wireless implant. By directing the stimulating leads through the blood vessels we can target specific nerves that are difficult to reach with traditional surgeries. Furthermore, we demonstrate this endovascular nerve stimulation (EVNS) with a millimeter sized wireless stimulator that can be delivered minimally invasively through a percutaneous catheter which would significantly lower the barrier to entry for neuromodulatory treatment approaches because of the reduced risk. This miniaturization is achieved by using magnetoelectric materials to efficiently deliver data and power through tissue to a digitally-programmable 0.8 mm^2^ CMOS system-on-a-chip. As a proof-of-principle we show wireless stimulation of peripheral nerve targets both directly and from within the blood vessels in rodent and porcine models. The wireless EVNS concept described here provides a path toward minimally invasive bioelectronics where mm-sized implants combined with endovascular stimulation enable access to a number of nerve targets without open surgery or implantation of battery-powered pulse generators.

---

Bioelectronic modulation of neural activity is a powerful tool for treating many disorders, especially when these disorders cannot be effectively managed with conventional therapies. For example, electronic devices that stimulate neural activity are effective for treating disorders like Parkinson’s Disease, epilepsy, chronic pain, hearing loss and paralysis [1–7]. These devices are most effective when implanted in the body where they can selectively stimulate the desired nerve targets; however, the invasiveness of the implantation can introduce additional risk for the patient. Invasive implants can also lead to complications such as chronic inflammation which can further degrade device functionality and lead to failure [8–10].

The vascular system that accompanies nerves as a part of the neurovascular bundle, provides a less invasive route for approaching nerve targets [11]. Existing neural implants for nerve targets such as the dorsal root ganglion (DRG) can suffer from site infection that results in device explantation and follow-up surgeries [12]. Endovascular neural stimulators (EVNS) delivered via an intravascular catheter to deep tissue targets with a minimally invasive procedure through the blood vessels within the body would leave the tissue target undisturbed. As a result, endovascular deployment of devices is often associated with significantly lower risk compared to open surgical approaches: recovery times are drastically reduced and site infections are extremely uncommon [11]. Given these advantages, an endovascular approach to neural stimulation would be attractive for the multitude of central and peripheral nerve targets that are adjacent to vascular structures, such as targets in deep brain, peripheral nerves, and the heart. [13–15]. Recently, several new endovascular bioelectronic devices have been developed that exemplify the benefits of stimulating neural tissue through the vasculature [16–19]. However, these devices have stimulation leads that are wired to pulse generators or centimeter-sized inductive coils. The long lead wires and implantation of centimeter-sized devices create additional failure points and require an open surgery that reduces some of the benefits of an endovascular surgical approach [20].

By miniaturizing the bioelectronic implants to a diameter of a few millimeters it would be possible to deliver endovascular neuromodulation therapies entirely with minimally invasive procedures that rely on percutaneous catheters. In order to sufficiently miniaturize the device to the size constraints of the catheter (< 3 mm in diameter), some form of wireless power is necessary to replace the bulkier batteries if we expect long-term operation. While several innovative wireless power transfer modalities have been demonstrated including far-field RF radiation, near-field inductive coupling, mid-field electromagnetics with hybrid inductive and radiative modes, ultrasound, and light; there has yet to be a demonstration of a millimeter-sized wireless neural stimulator that operates at a depth of several cm in a large animal model [21–34].

Here we turn to magnetoelectrics (ME) as a wireless data and power transfer technology due to its large power densities, high tolerance for misalignment, and ability to operate in deep tissue when compared to alternative wireless power technologies for bioelectronic implants [35].

Our results show for the first time that it is possible to safely stimulate peripheral nerves using electrodes placed inside the blood vessels, and that we can deliver the stimulation using a millimeter sized bioelectronic implant. By combining ME data and power delivery with a custom application specific integrated circuit (ASIC) we achieve a miniature device that is only 3 × 2.15 × 14.8 mm when fully encapsulated. Compared to miniature ultrasound powered devices, our MagnetoElectric-powered Bio ImplanT (ME-BIT) maintains functional power levels over a larger range of translational and angular misalignment, and does not need ultrasound gels or foams to couple energy from the transmitter. We also demonstrate a robust communication protocol that allows us to adjust the ME-BIT stimulation parameters. As a proof-of-concept, we show that these ME-BITs can be powered several centimeters below the tissue surface and can electrically stimulate peripheral nerve targets through the vasculature in a large animal model. These proof of principle studies open the door to minimally invasive bioelectronic therapies based on EVNS.

## Results

### Magnetoelectrics combined with a custom ASIC enables a millimeter-sized neural stimulator

To overcome the challenge of wireless data and power delivery to miniature bioelectronic implants, we developed a data and power delivery system based on magnetoelectrics, which achieves high power densities within the safety limits for human exposure [36]. Magnetoelectric materials provide efficient power delivery for bioelectronic implants by directly converting magnetic fields to electric fields based on the material properties [35, 37]. In our case we use a laminated bi-layer material that consists of Metglas, a magnetostrictive layer and lead zirconium titanate (PZT), a piezoelectric layer. When we apply a magnetic field to the material, the magnetostrictive material generates a strain that is coupled to the piezoelectric layer that, in turn, generates an electric field [35]. Thus, by applying an alternating magnetic field at the acoustic resonant frequency of the film we can efficiently deliver power to our implant [35,36,38, 39]. In addition to delivering power, we can also transmit data to our implant by modulating the frequency of the applied magnetic field. The frequency shift results in a change in the amplitude of the received voltage, which can be interpreted as a digital bit sequence that specifies the stimulation parameters for the implant [38,39]. Taken together, the complete wireless EVNS system consists of an external magnetic field transmitter, a ME film that harvests power and data from the magnetic field, and a custom integrated circuit that interprets the digital data and generates the electrical stimulus delivered by the electrodes (Fig. 1a). Figure 1b demonstrates a conceptual overview of the system implemented in a large animal model where a surface coil can be used to wirelessly transmit a magnetic field to power and program the implant for endovascular stimulation.

**Fig 1.**
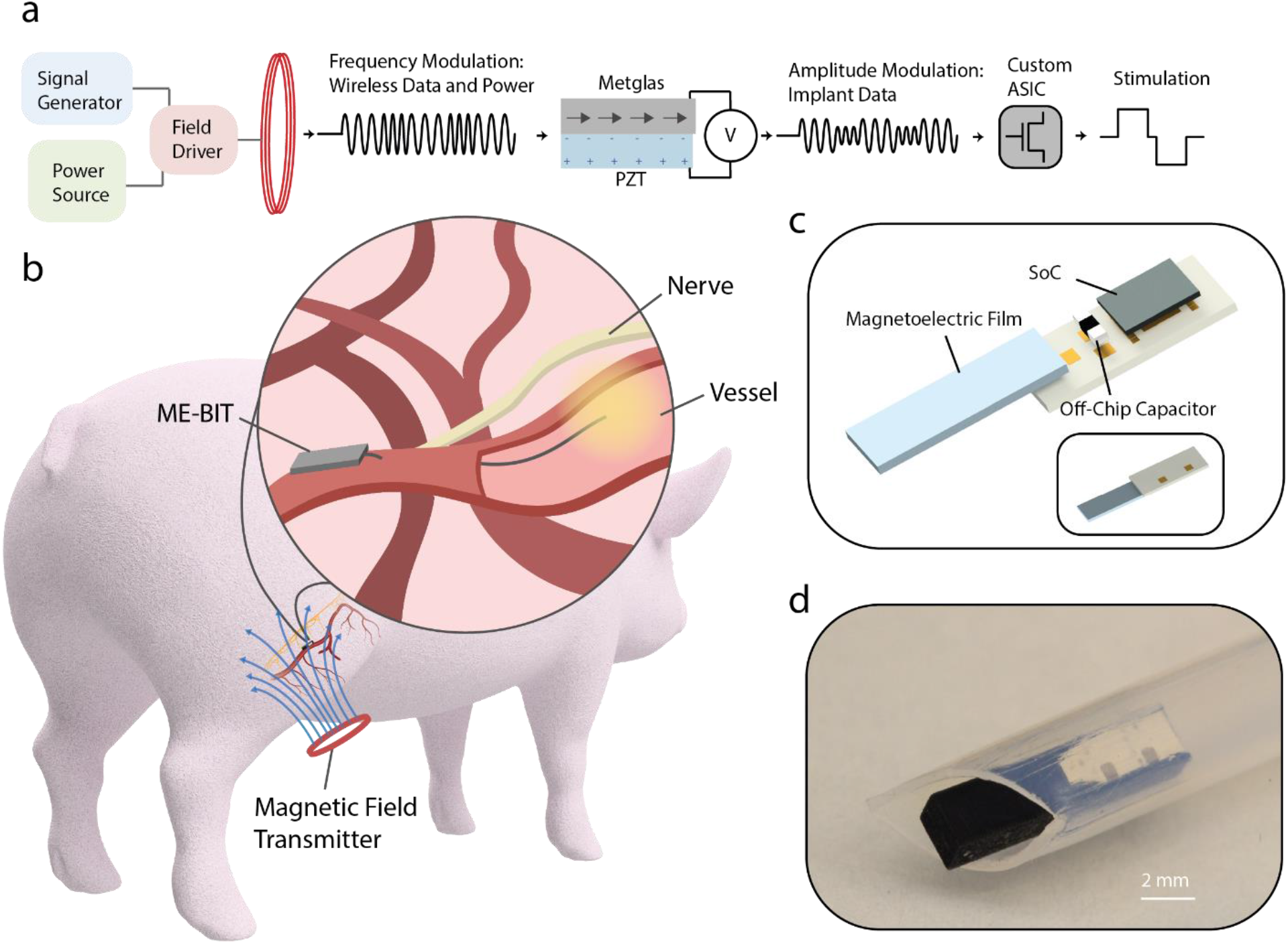
Wireless Magnetoelectric EVNS Overview: **a.** The overall EVNS system includes a magnetic field driver connected to a resonant coil that is tuned to the ME film resonant frequency. The coil outputs a frequency modulated magnetic field and switches between three different frequencies to modulate data and power received by the ME-BIT implant. The magnetic field is converted to an electric field by the magnetoelectric film. Specifically, the magnetostrictive layer, Metglas, mechanically deforms under the magnetic field and transfers the resulting strain to the piezoelectric layer, PZT, and induces a voltage. The amplitude of the voltage is then modulated by shifting the frequency of the applied field. The resulting voltage modulation received by the ME-BIT programs the custom IC to output the desired stimulus waveform. **b**. For our proof-of-concept experiments the ME-BIT is implanted proximally to a blood vessel deep within tissue and wirelessly powered through a magnetic coil in a pig. The implant’s stimulation lead is introduced into the vessel to stimulate nearby nerve targets. **c.** A rendering of the implant is shown with all the external components including the SoC, external capacitor, and the ME transducer. **d.** Photograph of the device fully packaged inside a 3D printed capsule resting in a clear sheath. The implant is encapsulated with a non-conductive epoxy prior to being implanted into the body and has the potential to be delivered endovascularly.

The ME-BIT itself consists of a magnetoelectric film with a size of 1.75 mm × 5 mm and a thickness of 0.3 mm for wireless power and data transfer, an ASIC for modulating the ME power and stimulation, and an external capacitor for energy storage as shown in the rendering in Fig 1c, which can be packaged to fit within an 11 French catheter. For our experiments we packaged the ME-BIT within a custom 3D printed PLA capsule with on-board electrodes that can also be used to power external electrodes. (Fig 1d) With this design, the miniature capsule can not only be delivered through a minimally invasive catheter, but also serve as a complete neuromodulatory device that can receive power, undergo programming, and transmit stimulation to neural tissue.

### A custom magnetic field transmitter enables data and power transfer at centimeter depths within safety limits

To deliver data and power to the implant we designed a magnetic field transmitter that drives a high-frequency biphasic current into a resonant coil [38]. By maintaining transmitter power levels below 1W, we can achieve field strengths of > 1mT sufficient to power the ME-BIT at depths of 4 cm within the safety limits.

Because the amplitude of the ME voltage peaks at the acoustic resonant frequency, we can send digital signals to our ME-BIT by detuning the applied magnetic field frequency. Figure 2 shows our communication protocol with charging, data transfer, and stimulation phases. As seen in Figure 2b we can select 3 frequencies to transmit digital data. The first frequency “Data 1” corresponds to the mechanical resonance (345 kHz). This is the frequency of maximum voltage (and maximum power transfer), which we use as a digital 1, and for the charging and stimulation phases. The second frequency “Data 0” is detuned by ~ 5 kHz. This frequency of 350 kHz produces a lower amplitude voltage, which is used as a digital 0. The third frequency is detuned by 55 kHz from the resonance peak and produces an even lower voltage than the “Data 0” signal. This “Notch Frequency” of 400 kHz is used to indicate the start of the data transfer and stimulation phases. By using the mechanical properties for the ME film to receive data based on frequency modulation we can avoid turning the transmitter coil on and off, which would require a settling time of 100 us for our resonant transmitters. Given the fast settling time of this frequency modulation scheme we find that 64 cycles of the carrier frequency can reliably transmit one bit, resulting in a 4.6-kbps data rate. We used a digital payload of 18-bits per stimulation, which accounts for a preamble and real-time calibration of the demodulation reference. This payload combined with the charging phase yields a maximum stimulation rate of 1 kHz, which is well within the range of typical neural stimulation applications [38].

**Fig 2.**
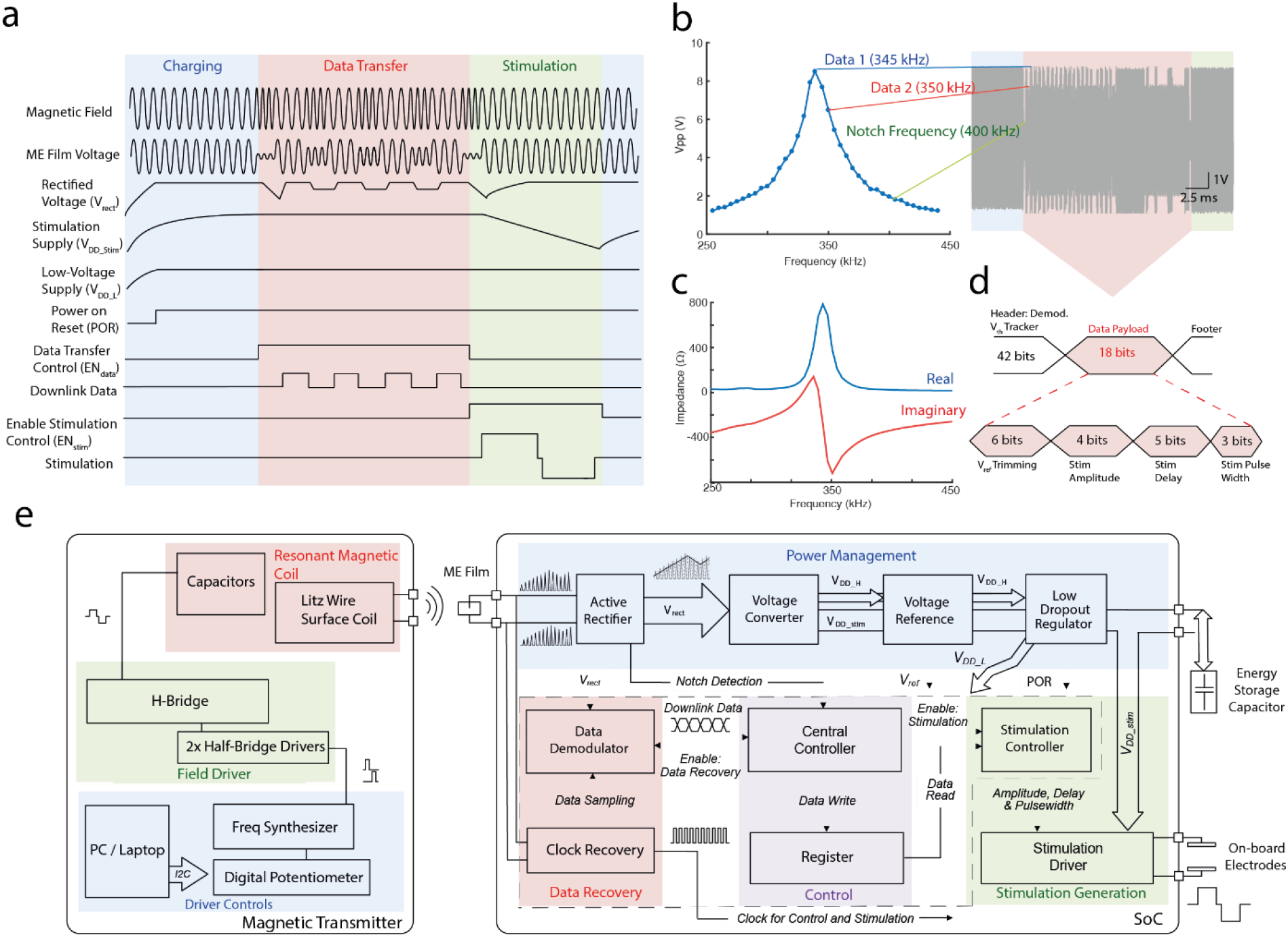
Timing and functional diagram for the ME-BIT. **a.** Timing diagram of the operation of the implant. The implant cycles through charging, data transfer and stimulating phase. The off-chip capacitor is charged in the charging phase to store energy for the stimulation. In data transfer, downlink data is received and demodulated by the ASIC to program stimuli. The operation of the implant is fully controlled by the transmitter through frequency modulation, which changes the carrier frequency of the magnetic field for different induced voltages of the ME film. **b.** The peak-to-peak voltage for a film resonating at 345 kHz as a function of magnetic field frequency. Three different field frequencies are chosen for communication with the SoC. Data 1 indicates the highest voltage which is used for the charging phase and for encoding “data 1”. Shifting the frequency by 5 kHz results in a 25% drop in Vpp that is used for “data 0” as can be seen in the data packet highlighted in red. The third frequency or notch frequency is chosen at a >50 kHz shift, which causes the Vpp to drop >85%. This is used to signify the beginning and end of the data transfer phase. **c.** The low film impedance at resonance lies at ~700 Ohms to support milliwatt level power budgets. **d.** A summary of the data payload including the header for demodulation threshold calibration and bits used for programming stimulation amplitude, delay, and pulse width. **e.** Block diagram of the magnetic transmitter (left) and the custom ASIC of the implant (right). The magnetic transmitter contains a controller, a magnetic field driver and a resonant magnetic coil to generate a low-frequency alternating magnetic field. The ASIC interfaces with a ME film to wirelessly receive power and data, and consists of power management, data recovery, control and stimulation modules to drive programmable stimulation. Energy for the high-power stimulation is stored in the off-chip capacitor, and stimulus is delivered through the on-board electrodes.

We estimate that this device can generate a maximum of 4 mW given the peak resonance voltage of > 8 Vpp with a resistive source impedance lower than 1 kohm (Fig 2d). This power level is sufficient for many neural stimulation applications [40].

### Custom Integrated Circuit provides digitally programmable stimulation with less than 9 uW power consumption

To deliver reliable stimulation independent of the coupling between the transmitter and the ME-BIT, the implant includes a custom ASIC that uses the digitally received data to program the shape (mono-phasic or bi-phasic), the amplitude (0.3 V to 3.3 V with 4-bit resolution), the pulse width (0.05 ms to 1.2 ms with 3-bit resolution), and the delay (0.01 ms to 0.8 ms) of the stimulation. The stimulation reference voltage is also programmed by the downlink data to generate a stimulation supply voltage 10% higher than the desired amplitude. As a result, the implant achieves > 90% stimulation efficiency ηstim (ηstim = stimulation amplitude /stimulation supply) for 1.5-to-3.3-V stimulation amplitude which helps to alleviate heating issues of the device in high-power stimulation.

The ASIC, fabricated on 180 nm complementary metal-oxide-semiconductor (CMOS) technology (TSMC), measures only 1 by 0.8 mm while performing several functions to ensure robust stimulation and communication. The ME induced alternating voltage is firstly rectified to the DC voltage Vrect by a full-bridge active rectifier with an 84% voltage conversion efficiency. This rectified voltage is then converted by an DC voltage converter, which provides proper voltage and buffers energy on the off-chip capacitor Cstore for stimulation. The voltage converter also generates a high-voltage supply VDD_H for other circuits for power management, including the low-dropout regulator (LDO) and the voltage references generator, and guarantees cold startup of the system as well. A constant supply VDD_L of 1 V is provided by the LDO for the controller, the data demodulator and the timing reference generator. To ensure proper system operation, a power-on-reset (POR) signal is triggered when VDD_L stabilizes.

To maintain reliable functionality of implants under varying ME voltages caused by change of transmitter-implant distance and alignment, the phase transitions of the IC are fully controlled by the transmitter through the short notches in ME voltage. In addition, the demodulation threshold for the amplitude-modulated data is generated autonomously in the beginning of the data transmission cycle to avoid the data recovery errors due to changes in ME voltage. Meanwhile, a global system clock is extracted from the source by a low-power comparator-based clock recovery circuit, ensuring process and voltage invariant timing references for data sampling and stimulation.

### Magnetoelectric implant demonstrates high tolerance towards misalignment

We find that our magnetoelectric-based power transfer approach displays improved tolerance for translational and angular misalignment when compared to other millimeter-sized implants. In comparison to inductive coils that harvest power based on magnetic flux, the ME materials harvest power based on magnetic field strength. As a result, it has been shown that ME demonstrates more stable power transfer as a function of angular misalignment. [36]. This angular stability is supplemented by the fact that the large magnetic permeability of the Metglas layer helps to concentrate the magnetic field lines along the length of the ME film [41].

Our simulations show that ME-BITs can tolerate approximately 3 cm misalignment from the center of the transmitter coil and a depth of 3 cm in tissue. Using COMSOL to model the magnetic field generated by our 15-turn transmitter coil we find an almost uniform magnetic field across 6 cm inner diameter of the coil (>70% of total transmitter area) as shown in Fig 3a. The dashed line in Fig. 3b shows the boundary line of 1 mT, which is the minimum operating field strength for the implant.

**Fig 3.**
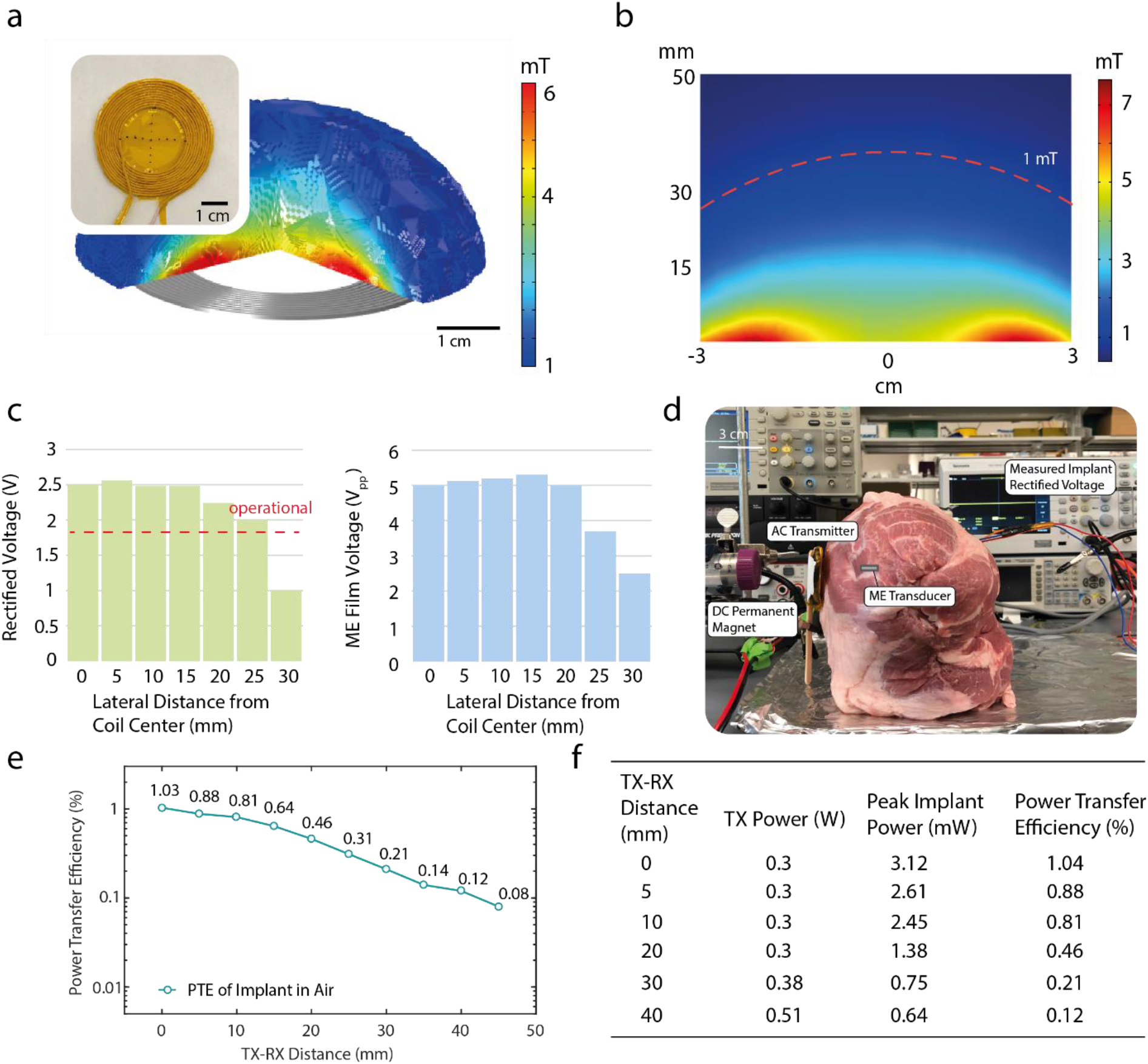
Characterization of ME power transfer. **a.** Finite element simulation of the magnetic field produced by one of the AC coil geometries used to power the ME implants. The coil shown is ~7cm in diameter made from 18 AWG litz wire and insulated with polyimide tape. The ME implant has been shown to maintain functional voltages at a field of 1 mT, and the COMSOL simulation demonstrates the space in which the implant remains functional where the edges are bound to be 1 mT. **b.** A 2-D cross section of the simulation from b is shown, where a 1 mT field can be generated at a distance of 30 mm and maintain lateral uniformity in case of translational misalignment. **c.** experimental data measuring the implant’s rectified voltage as the implant is moved away from the center of the coil. In order for the device to remain functional, the rectified voltage needs to be >1.8V as shown with the red dotted line. Also shown is the peak-to-peak ME voltage. **d.** An experimental image is shown for an ex-vivo model of porcine tissue. The magnetic field transmitter which is a combination of both the permanent magnet and AC coil, is placed on the left hand side of the tissue. The encapsulated ME film powering an implant is placed 2.5 mm within the heterogeneous tissue, while the rectified voltage of the implant is measured.**e.** Measured power transfer efficiency for the magnetoelectric implant as a function of distance in air. **f.** Table that shows transmitter power, peak implant power, and power transfer efficiency in air as the distance between transmitter and receiver is increased. Note that for this experiment, transmitter power was held constant until a distance of 2 cm, and then increased in order to maintain sufficient power to the implant.

When we tested our 15-turn transmitter coil we found that we could indeed power our ME films above our operating voltage (>3.6 Vpp) a distance of 3 cm in air from the surface of the coil with a misalignment tolerance that matched the 6 cm inner diameter of the coil. (Fig. 3c). This misalignment tolerance is more than 27 times greater than recently reported ultrasound-powered implants with a mm-scaled translational window [31]. The improved alignment tolerance would be advantageous for applications where an individual may want to align a transmitter several times a day or fit a wearable transmitter that may move and drift over time.

### Magnetoelectric power able to sustain operation of implant at centimeter distances within tissue

We found that the ME-BITs received enough power to function when implanted at centimeter depths in porcine tissue. (Fig. 3d) Specifically the ME film packaged inside the IC capsule was able to power the IC and maintain a sustained rectified voltage of 2.5V. By adjusting the magnetic field while increasing the distance between the implant and transmitter coil, the ME-BIT was able to maintain the optimal working voltage up to 2.5 cm. (Fig. S1) This energy was delivered through a ~3 mm air gap between the surface coil and tissue demonstrating the ability to achieve non-contact wireless power transfer.

When characterizing the power coupling efficiency in air we achieved functional power levels at a TX-RX distance of up to 4 cm, which is primarily limited by size and power level of our magnetic field transmitter. At the surface of the coil, we found that the ME-BIT generated a peak power of 3.12 mW resulting in a peak efficiency of 1.03%. (Fig. 3e) While we were able to keep input power constant up to 2 cm distances while still maintaining implant operation, the transmitter power was increased from 0.3 W to 0.51 W in order to keep a functional voltage on the implant at 4 cm. (Fig 3f) While the maximum distance demonstrated here is 4 cm, ME voltage is primarily dependent on magnetic field strength thus greater TX-RX distances may be achieved with optimization of driver electronics and transmitter designs.

### The ME-BIT demonstrates programmability and fully untethered operation for direct nerve stimulation in rat

Proof-of-concept experiments show that wirelessly powered ME-BITs evoke repeatable compound muscle action potentials (CMAPs) along with observable leg kicks when placed in contact with the sciatic nerve. This miniaturized implant had a volume of 6.2 mm^3^ and weight 30 mg which makes this suitable for small rodent models and was able to directly stimulate rat peripheral nerve (n = 2) in-vivo. (Fig. 1a) Stimulation for rat A while the mote was fully untethered and powered at a 1-cm-distance is shown in figure 1b, where a 3V, 1.5 mS pulse width monophasic pulse train is applied at 3 Hz. EMG recordings of foot muscles showed waveforms that were time locked with the applied stimulus at the same frequency. Often necessary in neural interfaces, the stimulation parameters on the ME-BIT can be adjusted by sending the appropriate commands through the magnetic field. Not only is the implant able to adjust its stimulation amplitude from 0.3V to 3.3V as shown in Figure 4c, it also is able to vary its pulse width and frequency to meet the demands of different neuromodulation applications and provide targeted therapies to account for variance from patient to patient. The programmability of the device is shown through an acute demonstration with rat B by varying the amplitude of the stimulus and observing a resulting graded EMG response. By adjusting the stimulation as well as the pulse width, the total charge delivered to the nerve could be controlled to directly affect the number of recruited motor units to elicit varying CMAP responses. In Fig. 4d, we innervated the sciatic nerve with monophasic pulses at 1 Hz while holding the pulse width at 1.5 ms and varying the amplitude from 300 mV to 3.1 V. The resulting CMAPs ranged in amplitude from 0.4 to 2.7 mV where the number of recruited muscle fibers seemed to saturate when increasing the stimulation amplitude from 2.1 to 3.1 V.

**Fig 4.**
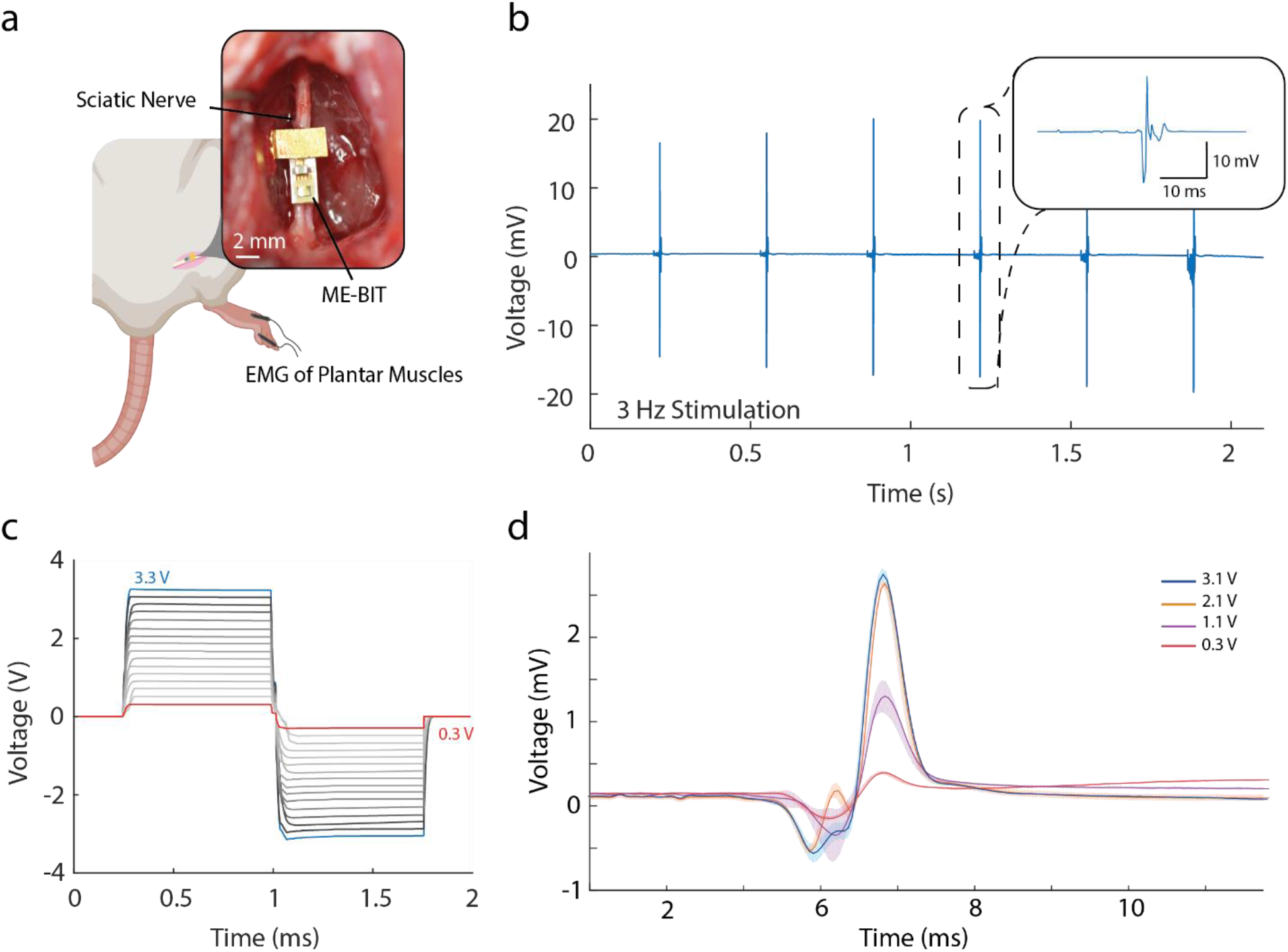
In Vivo Direct Nerve Stimulation in Rat Model: **a.** Schematic of the implantation of a fully wireless ME-BIT in a rat model. The device is placed on top of the rat sciatic nerve and wirelessly powered from an external transmitter. EMG recording electrodes are placed in the plantar muscles of the foot while a ground electrode is placed higher in the body. Also shown is an image of the actual device used for stimulation. **b.** The free-floating device is set to stimulate the nerve at 3 Hz with 3V, 1.5 mS pulses and the recorded EMG signal is shown. The insert shows a close-up of the highlighted EMG trace. **c.** A graph that shows programmed biphasic stimulus pulses of varying amplitudes. **d.** Averaged EMG recordings from the plantar muscles of the rat that show graded traces in response to varying levels of programmed stimulation powered by the wireless implant.

### The magnetoelectric system demonstrates wireless endovascular nerve stimulation for multiple nerve targets

To demonstrate endovascular neural stimulation and the potential for clinical translation, we implanted the ME-BIT in a pig and demonstrated peripheral nerve stimulation from within the blood vessels using a wirelessly powered device. For this experiment, the film was mounted along the PCB and soldered to gold pads with the stim lead wire soldered to an exposed pad on the top of the PCB prior to the device encapsulation (Fig 5a). For the surgery, an incision was made in the hind leg of the pig to expose both the femoral nerve and femoral artery. The ME implant is then placed into the surgical site and a 9 Fr sheath was then introduced into the femoral artery to allow access into the vessel. The parylene insulated stimulation wire connected to the implant is introduced into the vessel as shown in the schematic in Figure 5b. Images of the surgical site shows the femoral nerve and a sheath entering the femoral artery with the encapsulated implant placed proximal to the vessel. The magnetic field transmitter is then brought to the surface of the skin to wirelessly power the implant at an implanted distance of 1.5 cm. (Fig 5d) By applying a 3V monophasic stimulus pulse with 1.5 ms pulse width to the exposed tip of the endovascular wire, the device provided targeted monopolar stimulation with the reference electrode on the ME implant. As shown in Fig. 5c, we were able to stimulate the femoral nerve through the femoral artery at various stimulation frequencies including 10 Hz. Along with CMAPs, we recorded downstream nerve action potentials with bipolar hook electrodes shown in Fig 5e, as well as time averaged central somatosensory evoked potentials (SSEPs) (Fig. S3). These recordings demonstrate that the stimulation was mediated by the nerves and is not a direct muscle stimulation. To rule out that our data could be explained by stimulation artifacts due to the applied magnetic field, we performed control experiments with the magnetic field detuned from the ME resonance wavelength. (Fig. 5c) Although we transmit the same communication protocol, because the magnetic field is detuned, the ME-BIT does not accurately receive the digital data and the implant does not deliver a stimulus.

**Fig 5.**
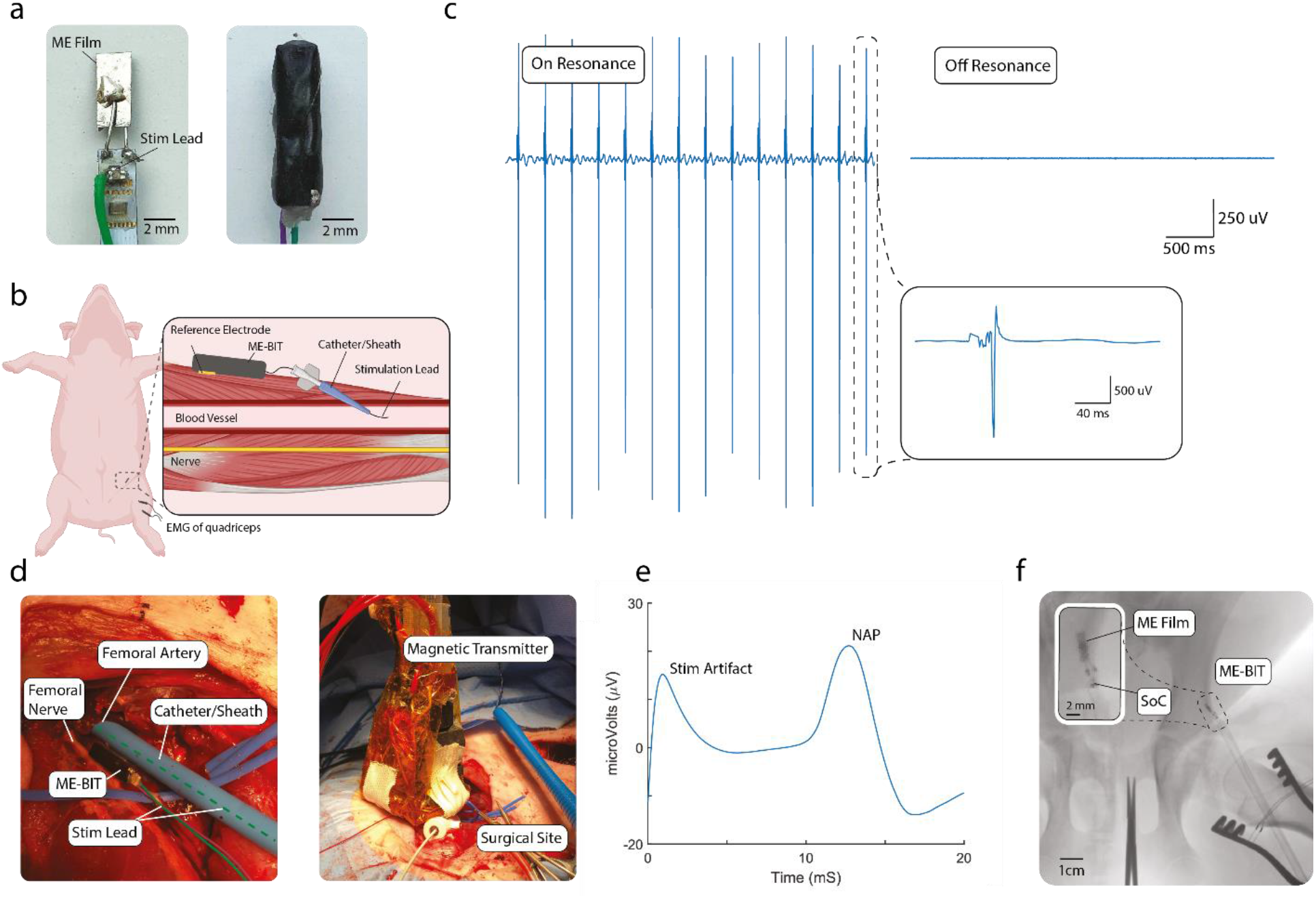
Endovascular Peripheral Nerve Stimulation in Large Animal Model: **a.** The exposed implant with ME film and soldered stimulation lead from the top of the printed circuit board (PCB). To the right is the fully encapsulated device in the 3D printed box and covered in non-conductive epoxy. **b.** A schematic for the endovascular stimulation in the pig. The implant is placed near the targeted femoral artery and the stimulation lead is introduced into the vessel through a catheter. **c.** The resulting EMG is shown where recording electrodes are placed on the bicep femoris of the pig. The blue trace on the left shows monophasic stimulation at 10 Hz with the magnetic field on resonant, while the trace on the right is the control where the magnetic field and transmitter are tuned to be off-resonant. **d.** The image on the left shows a close-up of the surgical site with a 9Fr sheath entering the femoral artery as well as the implant and femoral nerve. The image on the right shows the magnetic field transmitter on top of the skin powering the intravascular device and stimulating the femoral nerve at a distance of ~1.5 cm inside the body. **e.** A time averaged nerve action potential was recorded with bipolar hook electrodes directly on the femoral nerve with wireless endovascular stimulation. **f**. A x-ray where the ME implant is introduced into the femoral artery where the ME film, capacitor, and SoC can be seen.

We were also able to demonstrate EVNS of the intercostal nerves and the dorsal root ganglion, which are common targets for treating chronic pain [42]. As shown in Fig S2, we were able to introduce the insulated stimulation wire through the segmental artery to reach intercostal nerves and the DRG of the pig, which are both in direct contact with the intercostal arteries. Similarly to femoral nerve stimulation, we applied a monophasic stimulus pulse at 3V, 1.5 ms with varying frequencies from 1-10 Hz. The delivered exposed wire tip served as the monopolar electrode while the return electrode was an on-board electrode located on the ME implant. We also were able to measure compound muscle action potentials that resulted from the stimulation and observed time-aligned muscle contractions in the chest wall. When we performed the same off-resonant frequency controls as we did for the femoral nerve stimulation we found no response, supporting the fact that EVNS of the intercostal nerve was also a nerve stimulation from within the blood vessel.

The small mm-sized form factor of our ME-BITs also enables delivery of the entire device within the vasculature. As a proof of concept, we deployed our ME-BIT through an 9Fr sheath into the femoral artery as can be seen in Fig 5f. The ME film and ASIC is visible through X-ray and can allow for visualization and monitoring of the device post implantation. Tissue samples for histologic evaluation were also taken to assess for acute vascular damage from endovascular stimulation; no damage was observed as shown in Fig. S4–S6.

In summary, we demonstrate that not only do these miniature ME-BITs have sufficient power density to stimulate a clinically relevant large animal model from within a blood vessel, we also demonstrate the potential for wireless EVNS of multiple peripheral nerve targets. Utilizing the advantages of magnetoelectrics, the ME-BIT can be implanted deep within the tissue close to targeted areas without requiring lead wires that connect to a more superficial inductive coil.

## Discussion

This work represents the first example of a magnetoelectric-powered bioelectronic implant in a large animal model, and highlights several of the advantages of this wireless power transfer technology for biomedical applications. Specifically, it’s large angle and lateral misalignment tolerance are favorable for the future use of wearable transmitters to power and communicate with the ME-BIT. While the implant itself might only move a few millimeters once fixed within the tissue, it is easy to imagine misaligning a wearable transmitter by a centimeter or more, which remains within our alignment tolerances. Furthermore, the use of a wearable transmitter is also possible due to the low magnetic field strengths that are required to activate high voltages in these ME thin films where only ~1 mT field strengths are required for the power densities required to activate neurons through stimulation by the material itself [35] or by powering custom integrated circuits [38,39]. This will allow the technology to be readily translated into the clinic and even permit patients to use implants at a home-setting. Furthermore, because magnetoelectrics have excellent scaling properties for wireless power transfer compared to other methods [33], it may be possible to significantly reduce the size of the device to the point where it could fit in smaller vessels and be deployed to difficult to reach targets.

The data presented here shows the first proof-of-concept that peripheral nerves can be stimulated from within the blood vessels using a millimeter-sized wireless implant. Additional work is needed to develop this technology into a biomedical device for clinical use. For one, hermetically sealed packaging will be needed for chronic implantation of the device. While thin film packaging solutions have yet to be fully developed for clinical use, other wireless implants have shown that glass, ceramic, or metal (Ti) casings can enable chronic operation [43]. Fortunately, the magnetic fields should easily penetrate these materials and thus they are not expected to degrade the power coupling efficiency. Long-term deployment of future endovascular bioelectronics may also require adjunctive therapies using blood thinning pharmaceuticals. Several factors that involve the implantation of devices within the vasculature can promote thrombosis; however, improved techniques along with antithrombotic regimens have been shown to decrease any catastrophic thrombosis due to stent implantations to <1% [44,45]. Furthermore, it has also been shown that extended implantation of cardiac pacing leads that develop occlusions can result in collateral venous channels that would reroute blood around the occlusion [46]. Future studies would be warranted to determine how chronic deployment of the ME-BIT within the blood vessel could affect vasculature health.

Endovascular bioelectronics like the ME-BIT demonstrated here, opens the door for a wide variety of therapies that involve low risk and high precision implantable devices. Having bioelectronics implanted within the vasculature enables devices to be implanted in many parts of the body that are traditionally difficult to reach without having major risks for surgery. Additionally, bioelectronic implants with access to the bloodstream could enable real-time sensing of biochemicals, pH, or oxygenation levels within the blood to provide diagnostics or support closed-loop-electronic medicine [47,48]. Overall, wirelessly powered millimeter-sized devices implanted within or near the vasculature opens up numerous opportunities for minimally invasive bioelectronic medicine.

## Methods

### Fabrication of ME-BITs

The external capacitor was mounted on the PCB before wirebonding the custom IC to the board. After wirebonding the die area is encapsulated with epoxy to maintain the structural integrity of the bonds. The magnetoelectric film is fabricated with a 127 um thick PZT (APC Int.) bonded to a 23 um thick layer of Metglas (Metglas Inc.) with a thin epoxy layer (Hardman Double/Bubble). The films are then laser cut by a femtosecond laser cutter to the desired shape. Depending on the iteration of the device, the geometry and strategy for interfacing the film with the board is slightly different. For the direct nerve stimulator, the film size was manufactured to be ~4 mm × 3 mm. This film was then coated with ~10 nm of Ti and ~40 nm of Au with RF sputtering. The film’s non-coated side is bonded to an exposed pad on the PCB with conductive Ag epoxy (Electron Microscopy Sciences) and allowed to cure at 60 C for 20 minutes. The top electrical connection of the film is made by wirebonding the Au coated side to the second exposed pad on the PCB. In the case of the endovascular device, the aspect ratio is more important as the device length is not as important as the width in order to fit inside a catheter/sheath and deployed in blood vessels. The films are cut out to be ~1.75mm × 5mm. Conductive Ag epoxy is used to connect 30 AWGwire to the center of the ME film. This ME film is then soldered directly to the two exposed pads on the PCB. The insulated stimulation wire is then soldered onto the top side of the PCB. A second wire shown in Fig. 5a and 5d (purple) is soldered to the reference electrode for more flexible positioning but was ultimately not used in the experiment. Since the endovascular stimulator is introduced into the animal and can be placed within the blood vessel, a 3D printed capsule is designed and printed from PLA. The assembled implant is placed within the capsule with final dimensions of 3 × 2.15 × 14.8 mm and sealed with non-conductive epoxy.

### Device Testing and Calibration

Before fully packaging with an enclosure for encapsulation, each device was checked through comprehensive functional tests. In addition to the pads connected to the film, the energy storage capacitor and the stimulating electrodes, the ASIC also has testing pads and readout circuits providing gateways to the internal signals, such as the verified voltage, the LDO output, and the demodulated downlink data. Through monitoring these signals at various conditions, like different TX-RX distances and misalignments, we validated that the devices can operate properly in the future implantation. The ASIC was designed with robustness against source amplitude changes and process variations, in addition to this, we calibrated some variables during the tests to further improve the reliability of device operation and the effectiveness of stimulation. First, the carrier frequency shift for amplitude modulation needs to be carefully set. Simply employing a large enough frequency change may ensure that the voltage difference between data “1” and data “0” is always sufficient (> 100 mV), but would sacrifice the received voltage and power in the data transfer phase and hence suffer a smaller TX-implant distance. Therefore, an optimal frequency setting needs to be found to maximize the ME induced voltage while still providing a large enough modulation index for correct data demodulation. Due to the process variations in manufacture of the ME laminate, this optimal frequency shift may demonstrate slight variabilities among different ME films. Second, the reference voltage for the stimulation driver is generated on-chip, as a result, its accuracy may be affected by the process variations in semiconductor fabrication. To assure effective stimulation, we could calibrate the voltage reference generator with the downlink data to guarantee desired stimulating voltages at any cases.

### ME power transfer characterization

Ex-vivo porcine tissue that consisted of pork chuck, purchased from a local store, was a general mixture of mostly muscle tissue mixed with some fat and connective tissue. The 15-turn magnetic coil was held in place vertically with a small ~3 mm air gap from the piece of tissue and the permanent bias magnet was placed ~2 cm away from the AC coil. In order to maintain the feedback pins on the device, the ME-film was encapsulated in the endovascular 3D printed capsule and implanted with a stiffener into the distal end of the ex-vivo tissue and pushed through until the device reached the proximal end with the coil. The film was then used to power the ME-BIT and the various feedback signals were monitored and rectified voltage was recorded up to 2.5 cm depth within the tissue. To measure power transfer efficiency, we measured both the transmitter power as well as the peak implant power. Using a tektronix current probe (CT2 AC Current Probe) to measure current running through the AC coils as well as a potentiostat (Gamry Reference 600+ Potentiostat) to measure the impedance of the coil at the operating frequency, the transmitter power was calculated. The peak implant power on the other hand was measured by observing the charge current on the device powered by a 12mm^2^ ME film, where 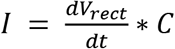, where C (~800 pF) is the on chip capacitor, and 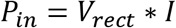

### Magnetic Field Transmitter

A microcontroller or computer that is capable of communicating via I2C protocol sends commands to a frequency synthesizer IC. The frequency synthesizer IC provides the three different data and notch frequencies used to communicate with the custom IC. This signal then is modulated through various circuits before being transmitted to half bridge drivers that are then passed to an H-bridge made up of 4 integrated driver ICs and MOSFETS (Texas Instruments CSD95378BQ5M). The H-bridge is rated up to 30 Amps continuous current with a upper frequency limit of 1.25 MHz.The data and notch frequencies are set by the user through a serial interface in Arduino as well as all of the stimulation parameters. The H-bridge is then connected to surface coils that are wrapped with 18 AWG Litz wire (MWS Wire) and resonated with high voltage rated capacitors (~6kV, WIMA). The field profiles were simulated using finite element modeling (COMSOL) and experimentally measured with an AC magnetic field probe (AMF Life Systems). The field profile was created by simulating a current of 18A in the 15 turn coil. Another magnetic field driver was also used for the porcine experiments. [49]

### In-Vivo Rat Stimulation Model

All procedures complied with the National Institutes of Health standards and were approved by the Animal Care and Use Committee of Rice University. (Protocol# IACUC-20-181) In-vivo stimulation with ME wireless power was confirmed in 3 different male Long-Evans rats (Charles River) ranging from 300g-400g. The stimulation was verified with a visually observed leg kick and corresponding EMG recording in the subplantar region of the foot. For the acute procedure, the animal was placed in an induction chamber with 5% isofluorane in oxygen at a flow rate of 1-2 liters per minute until the rat was unconscious and areflexic, confirmed with toe pinches. The animal was then transferred to a 40C heated pad with a nose cone with ~2% isofluorane. Meloxicam 2mg/kg SC and Ethiqa XR SC 0.65 mg/kg were administered to the rat before shaving the surgical site. Iodine swabs were used to sterilize the site before a single semi-circular incision was made across the lower hip of the rat. The fascial plane between the gluteus maximus and the anterior head of the bicep femoris was opened to expose the sciatic nerve. The underlying connective tissue was severed to better isolate the sciatic nerve.

Two EMG electrode needles were placed in the plantar muscles of the rat leg while a third ground electrode was placed on the main body of the rat. The recording electrodes were connected to a dual bioamplifier (ADsystems) and sampled at 1 kHz. The data is acquired and exported through Labchart and processed in Matlab 2017. On completion of the study, the animals were immediately euthanized under proper guidelines.

### In-Vivo Porcine Model

The animal procedures were conducted in accordance with the rules of the Institutional Animal Care (ANC) and Use Committee (Protocol# 2007074). 8 female Yorkshire pigs, weighing approximately 35–45 kg, received a 7-day acclimation period prior to any procedure. General anesthesia was administered by veterinary services personnel and was established with Telazol 4.4 mg/kg, ketamine 2.2 mg/kg, and xylazine 2.2 mg/kg IM, followed by intubation under general anesthesia. Mechanical ventilation was given with a mixture of oxygen and isoflurane (1–3%). Routine physiological monitoring was performed.

The pigs were placed in supine position and the femoral artery was palpated between the rectus femoris and the vastas medials muscles. A 6 cm skin incision was performed to expose the femoral neurovascular bundle and blunt dissection was used to remove the surrounding connective tissue and adventitia further exposing the femoral artery, vein, and nerve. Baseline EMGs (recorded from the quadricep muscles) and NAPs (recorded from the femoral nerve) were obtained from direct femoral nerve stimulation to assure nerve integrity after exposure. Swine analogues to Human 10/20 electrode positions were placed; Nasion (Ns), Cervical spine rostral (CSr), and Cervical spine caudal (CSd).

A 9 French sheath was then delivered into the common femoral artery (CFA) through a modified Seldinger technique. A parylene insulated wire (0.008-in) connected to the ME implant is introduced into the vessel. NAPs were then recorded on the femoral nerve and EMGs on the adjacent quadricep muscles after endovascular femoral nerve stimulation. Leg twitching was observed with each EMG recording. Central signals were also recorded from the cranial and cervical electrodes. Next, under direct fluoroscopic visualization, a 5-Fr Mikelson catheter (Cook Medical, Bloomington, IN, USA), was advanced over an 0.035-in Glidewire (Terumo IS, Somerset, NJ, USA) into the descending aorta at the level of a segmental artery. The same parylene insulated wire connected to the ME implant was then introduced into the segmental artery through a microcatheter (0.017-in inner diameter). Endovascular DRG and intercostal nerve stimulation was then performed through wireless magnetoelectric stimulation of the microwire. EMG was recorded from the intercostal muscles and twitches were seen with each recording. On completion of the study the animals were immediately euthanized under proper guidelines. The femoral and intercostal arteries tested were harvested for histopathologic examination. All tissue samples were routinely fixed in 10% formalin, processed and embedded in paraffin. Tissue blocks were sectioned at 5 um thickness for hematoxylin and eosin (H&E) and modified Movat pentachrome histochemical staining [50,51].

The electrophysiologic recordings were done on a Cadwell IOMax utilizing subdermal needle electrodes for recording for SSEP and EMG recordings. Direct nerve stimulation was performed via a triple hook electrode and direct nerve recordings were done via a double hook probe. Central response recordings were done via one subdermal needle placed over the rostrum referenced to a subdermal needle placed over the midline of the cervical spine. In some cases, a second subdermal needle was placed over the cervical spine with one positioned just behind the occiput and one placed at the lower cervical spine, always over the midline. A ground electrode was placed in the shoulder to help eliminate any unwanted artifacts. Amplification for central recordings and nerve action potentials was 100 uV/div and for all EMG recordings was 1000 uV/Div. Digital filter bandpass settings were applied for each type of recording: Central sensor responses (30-500 Hz), EMG recordings (10-3000 Hz), and nerve action potentials (30-1000 Hz).

## Supplemental

**Supplemental Fig 1.**
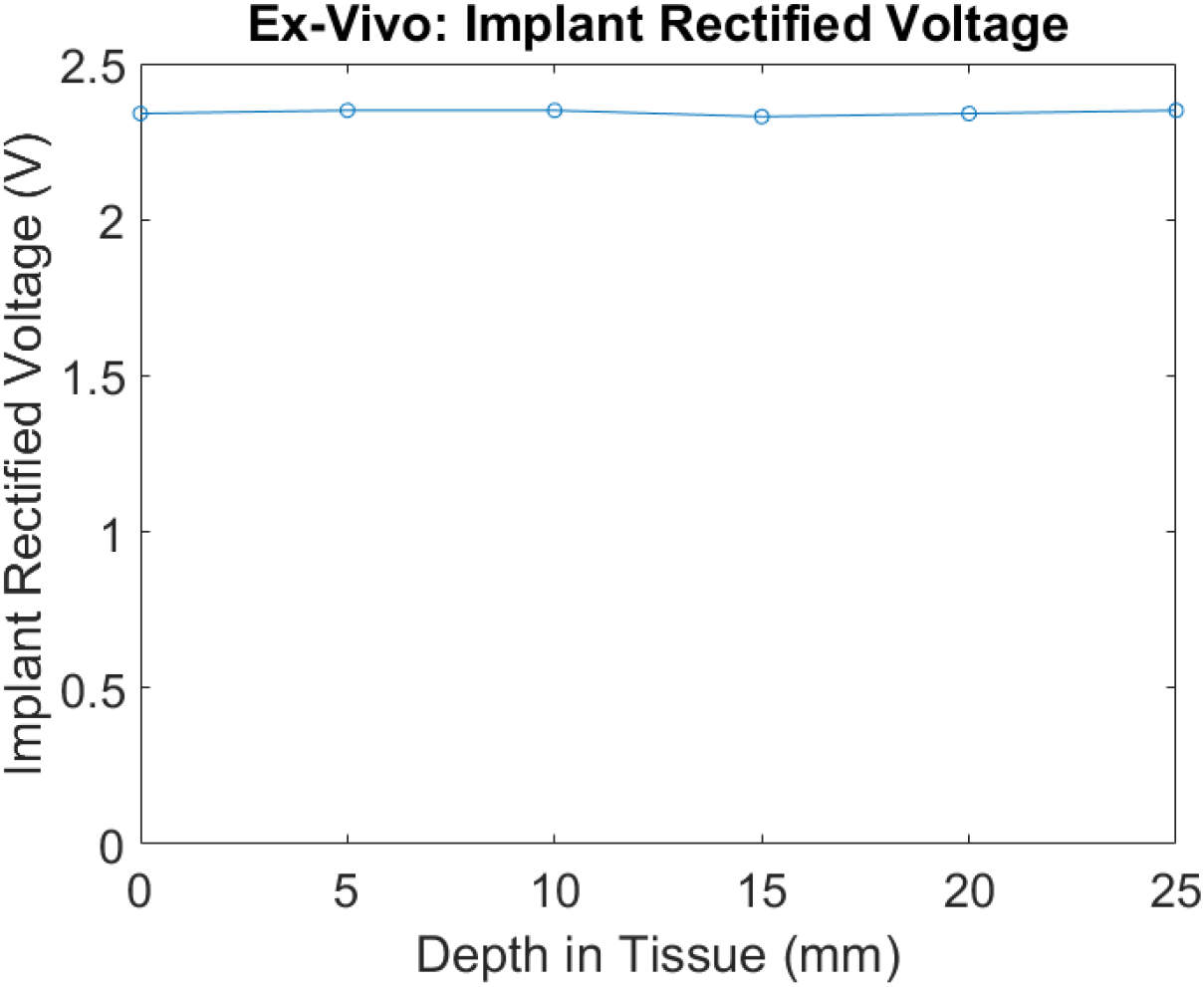
Rectified voltage of ME-BIT in ex-vivo porcine tissue while increasing TX-RX distance

**Supplemental Fig 2.**
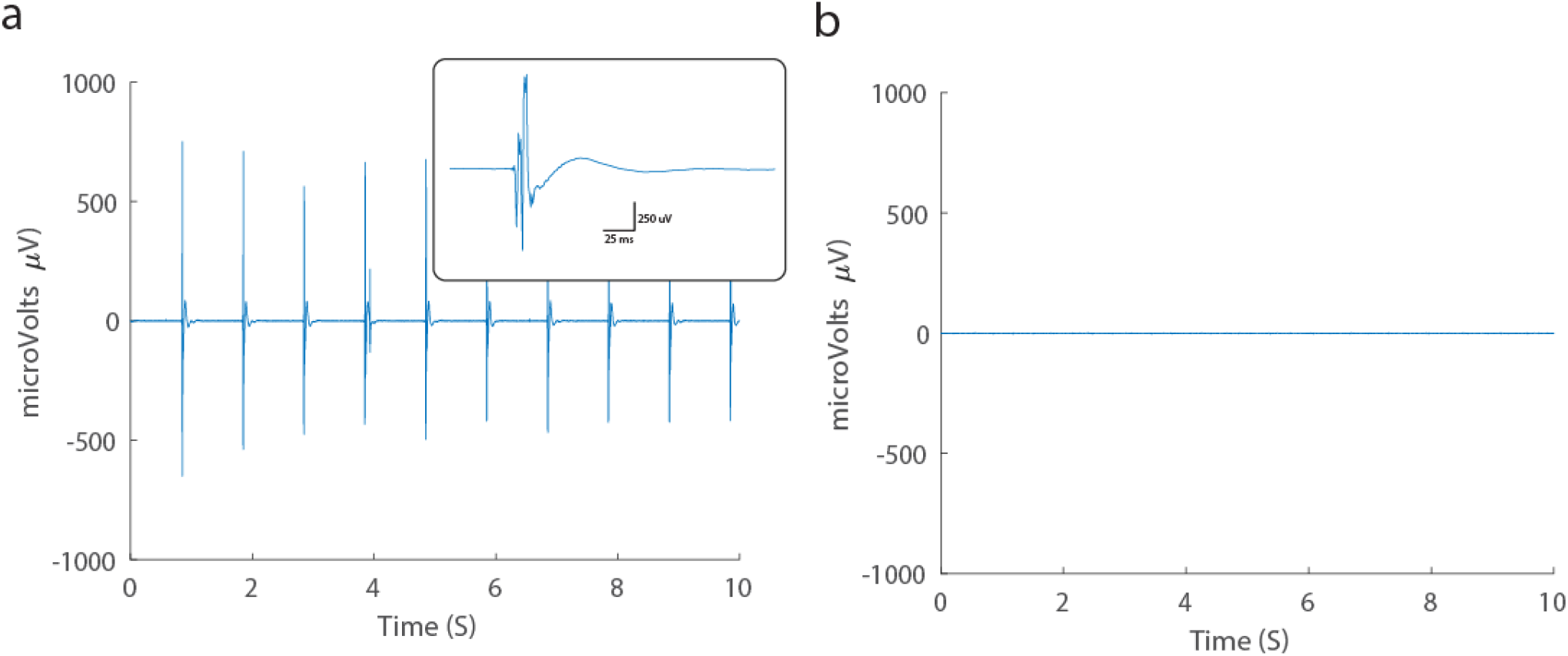
EMG recording for intercostal/DRG nerve stimulation (left) EMG recording of control by using off-resonant magnetic field.

**Supplemental Fig 3.**
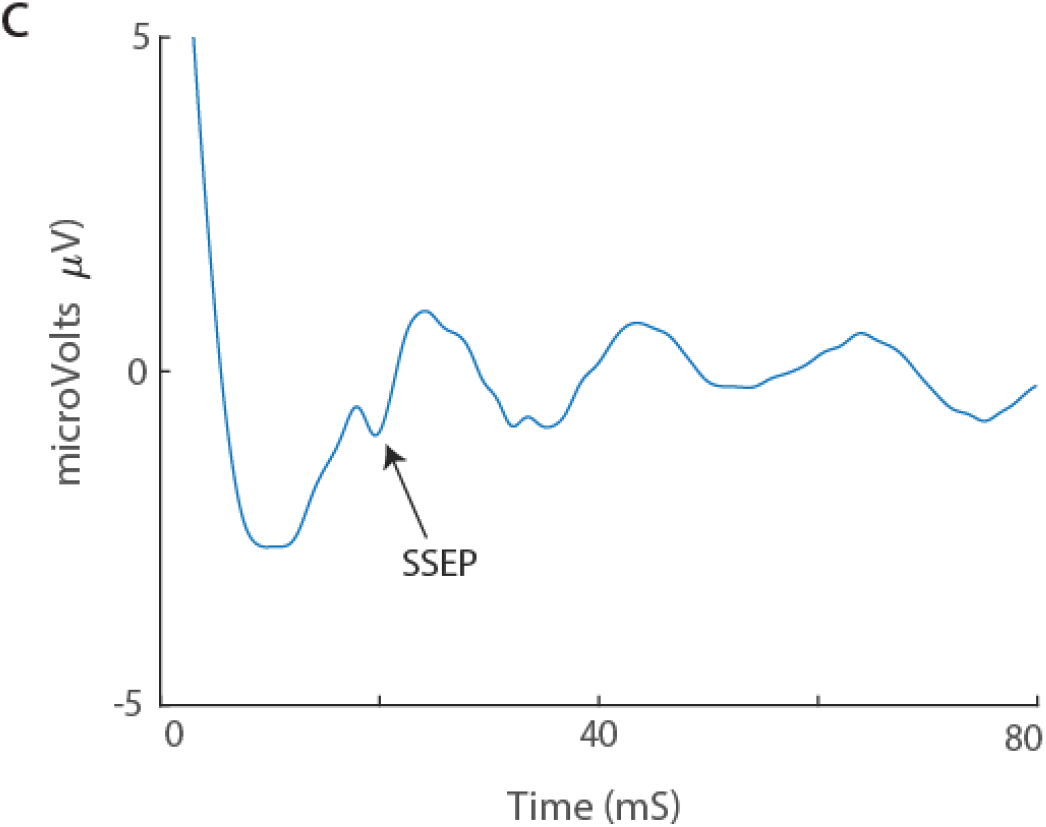
SSEP Central recording of the brain stem in response to femoral nerve stimulation

**Supplemental Fig 4.**
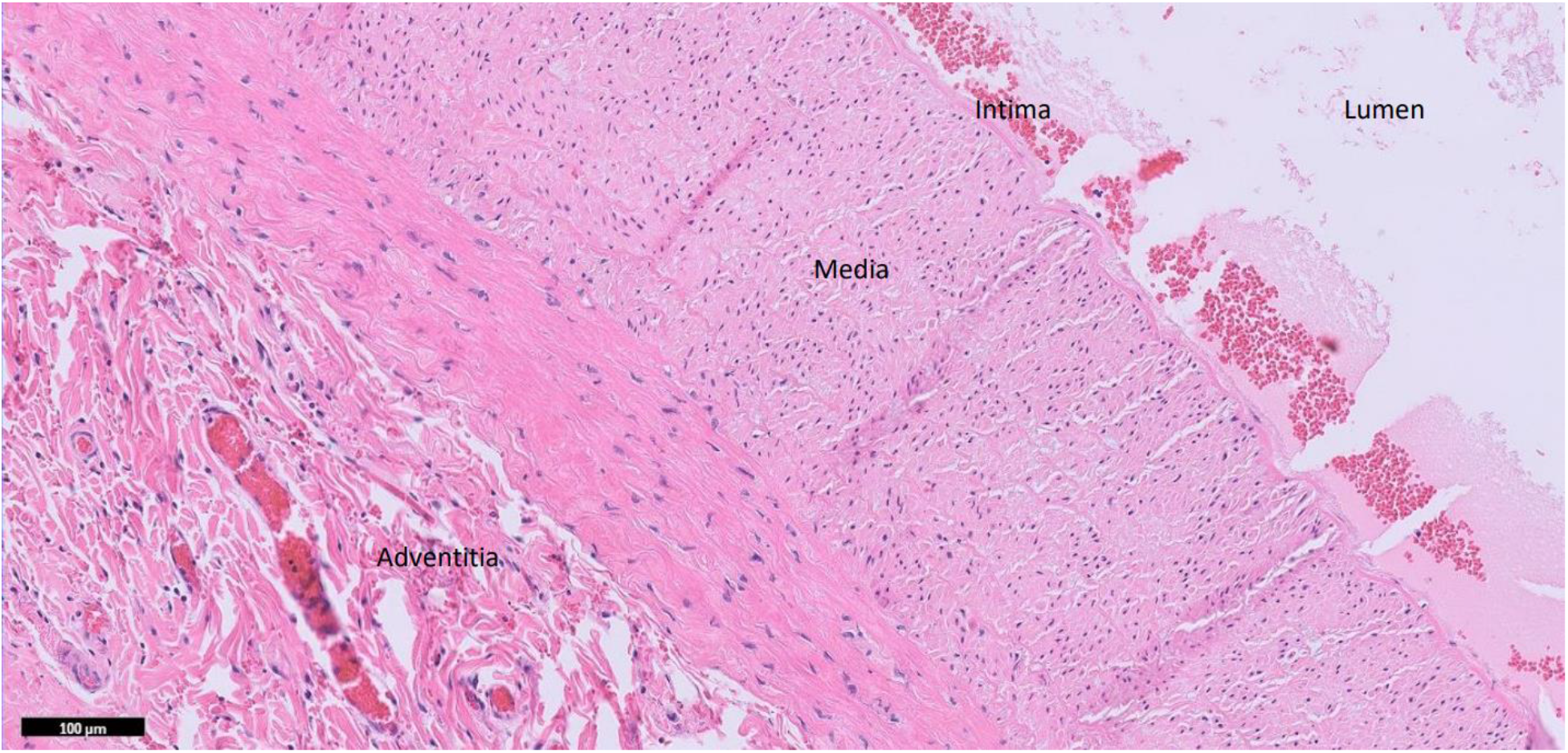
H&E stain of femoral artery which shows no evidence of acute vascular damage.

**Supplemental Fig 5.**
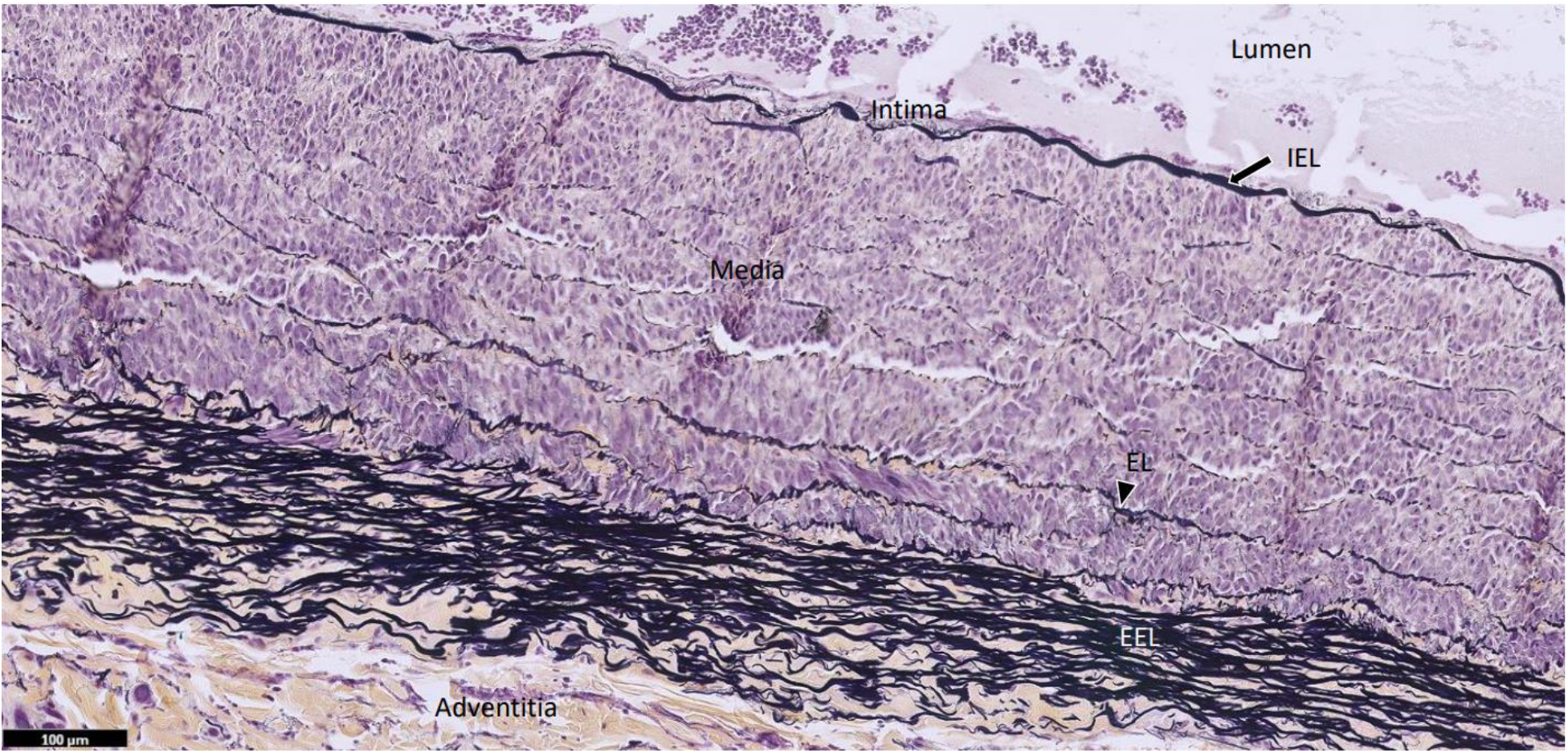
Modified Movat stain of femoral artery which also shows no evidence of acute vascular damage; Movat stain highlights normal connective and elastic tissue of the artery. (IEL = internal elastic lamina; EL = elastic lamellae; EEL= external elastic lamina)

**Supplemental Fig. 6.**
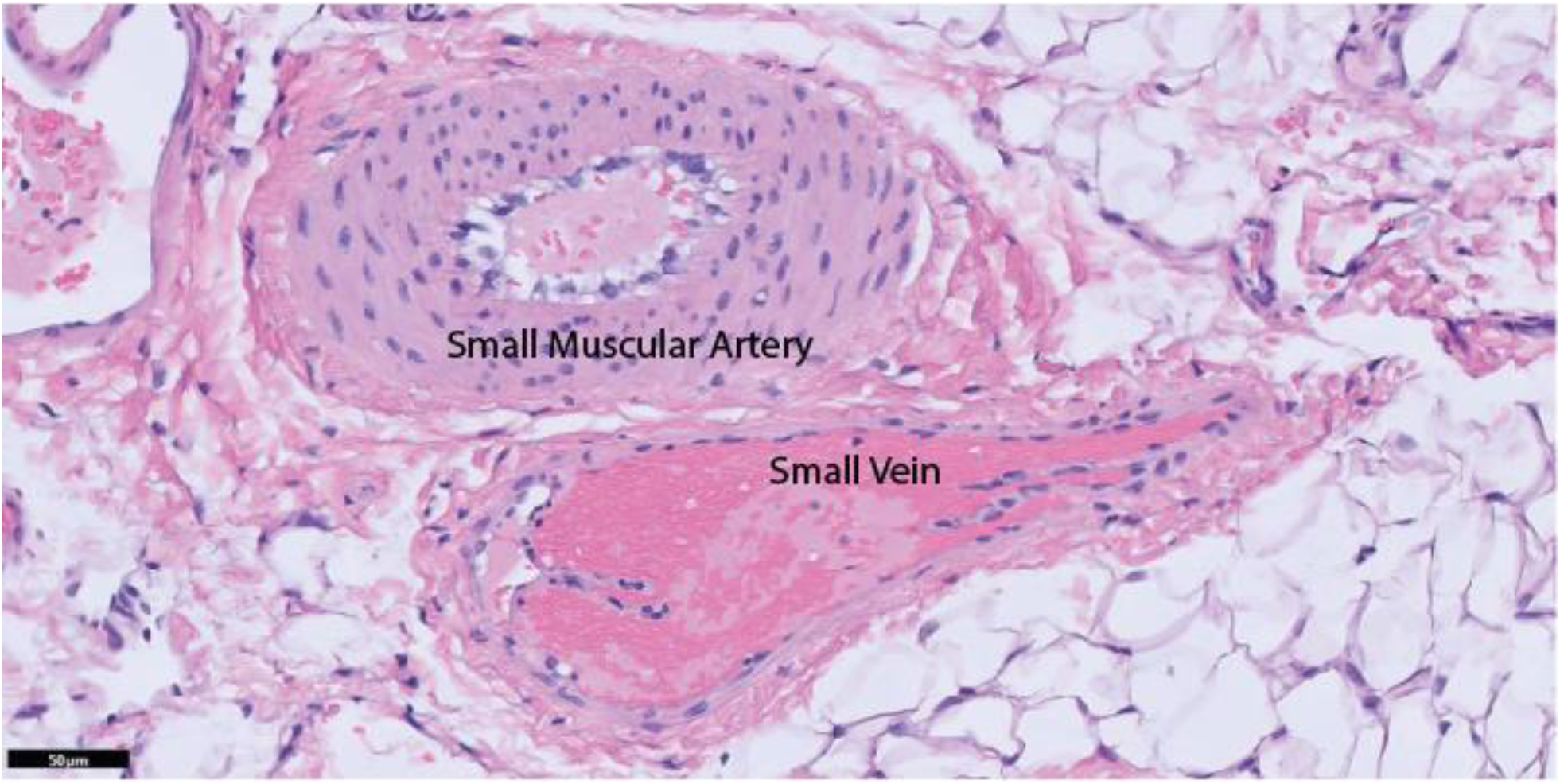
H&E stain demonstrates normal histology of the small vascular branches of an intercostal neurovascular bundle.

**Supplemental Figure 7:**
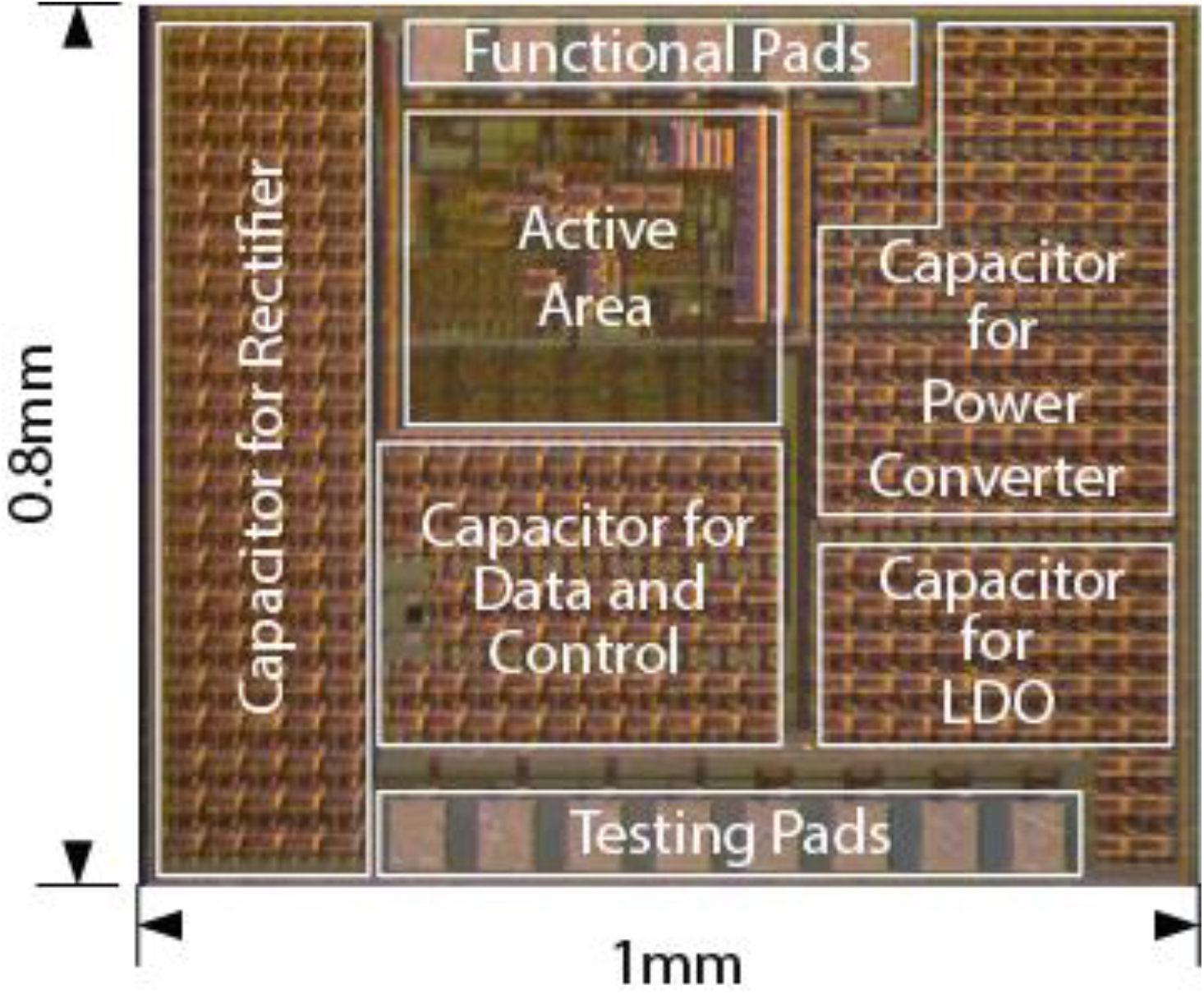
Custom ASIC Footprint and Layout

## Acknowledgements

This work was supported in part by the National Institutes of Health NIH grant no. U18EB029353 and R01DE021798; National Science Foundation GRFP (to J.C.C. and C.E.L.) The authors thank Boshuo Wang, Zhongxi Li, Angel Peterchev, and Stefan Goetz from Duke Univeristy for providing driver electronics for the porcine experiments.

## References

1. Shealy, C. N. Mortimer, J.T. and Reswick, J. B.. Electrical Inhibition of Pain by Stimulation of the Dorsal Columns: Preliminary Clinical Report. Anesthesia & Analgesia, vol. 46, pp. 489–491 (1967)

2. Verrills, P. Sinclair, C. and Barnard, A. A review of spinal cord stimulation systems for chronic pain. J Pain Res, vol. 9, pp. 481–492, (2016)

3. Morrell, M.J. Responsive cortical stimulation for the treatment of medically intractable partial epilepsy. Neurology 77, 1295–1304 (2011).

4. Cook, M.J. et al. Prediction of seizure likelihood with a long-term, implanted seizure advisory system in patients with drug-resistant epilepsy: a first-in-man study. Lancet Neurol. 12, 563–571 (2013).

5. Benabid, A. L. Deep brain stimulation for Parkinson’s disease. Current Opinion in Neurobiology, 13(6), 696–706. (2003)

6. Wilson, B.S. et al. Better speech recognition with cochlear implants. Nature 352, 236–238 (1991).

7. Hochberg, L.R. et al. Reach and grasp by people with tetraplegia using a neurally controlled robotic arm. Nature 485, 372–375 (2012).

8. Polikov, V.S., Tresco, P.A. & Reichert, W.M. Response of brain tissue to chronically implanted neural electrodes. J. Neurosci. Methods 148, 1–18 (2005).

9. McConnell, G. C., Rees, H. D., Levey, A. I., Gutekunst, C. A., Gross, R. E., & Bellamkonda, R. V. Implanted neural electrodes cause chronic, local inflammation that is correlated with local neurodegeneration. Journal of Neural Engineering, 6(5). (2009)

10. Butson, C. R., Maks, C. B. & McIntyre, C. C. Sources and effects of electrode impedance during deep brain stimulation. Clin. Neurophysiol. 117, 447–454 (2006).

11. Fan, J. Z., Lopez-Rivera, V., & Sheth, S. A. Over the Horizon: The Present and Future of Endovascular Neural Recording and Stimulation. Frontiers in Neuroscience, 14(May), 1–5. (2020)

12. Moman, R. N. et al. Infectious Complications of Dorsal Root Ganglion Stimulation: A Systematic Review and Pooled Analysis of Incidence. Neuromodulation: Technology at the Neural Interface (2021)

13. Penn, R. D., Hilal, S. K., Michelsen, W. J., Goldensohn, E. S., and Driller, J.. Intravascular intracranial EEG recording. J. Neurosurg. 38, 239–243. (1973)

14. Rush A. J. et al. Vagus nerve stimulation (VNS) for treatment-resistant depressions: a multicenter study. Biol. Psychiat. 47 276–286. (2000)

15. Oxley, T. J. et al. Minimally invasive endovascular stent-electrode array for high-fidelity, chronic recordings of cortical neural activity. Nature Biotechnology, 34(3), 320–327. (2016)

16. Opie, N. L., et al. Focal stimulation of the sheep motor cortex with a chronically implanted minimally invasive electrode array mounted on an endovascular stent. Nature Biomedical Engineering, (2018)

17. Scherlag, B. J. et al. Endovascular stimulation within the left pulmonary artery to induce slowing of heart rate and paroxysmal atrial fibrillation. Cardiovascular Research, 54(2), 470–475. (2002)

18. Oxley, T. et al. Motor neuroprosthesis implanted with neurointerventional surgery improves capacity for activities of daily living tasks in severe paralysis: First in-human experience. Journal of NeuroInterventional Surgery, 13(2), 102–108. (2021)

19. Hasdemir, C., Scherlag, B. J., Yamanashi, W. S., Lazzara, R., & Jackman, W. M. Endovascular stimulation of automatic neural elements in the superior vena cava using a flexible loop catheter. Japanese Heart Journal, 44(3), 417–427. (2003)

20. Sahin, M., & Pikov, V. Wireless microstimulators for neural prosthetics. Critical Reviews in Biomedical Engineering, 39(1), 63–77. (2011)

21. Montgomery, K. L. et al., Wirelessly powered, fully internal optogenetics for brain, spinal and peripheral circuits in mice, Nature Methods, vol. 12, pp. 969–974, (2015)

22. Loeb, G. Peck, R. Moore, W. Hood, K. BION System for Distributed Neural Prosthetic Interfaces, Medical Engineering & Physics, vol. 23, pp 9–18 (2001)

23. Shin, G. et al., Flexible Near-Field Wireless Optoelectronics as Subdermal Implants for Broad Applications in Optogenetics, Neuron, vol. 93, pp. 509–521.e3, (2017)

24. Freeman, D. K. et al., A Sub-millimeter, Inductively Powered Neural Stimulator, Front. Neurosci., vol. 11, (2017)

25. Jia, Y. et al., A mm-sized free-floating wirelessly powered implantable optical stimulating system-on-a-chip. 2018 IEEE International Solid - State Circuits Conference - (ISSCC), pp. 468–470 (2018)

26. Khalifa, A. et al. The Microbead: A 0.009 mm3 Implantable Wireless Neural Stimulator, IEEE Transactions on Biomedical Circuits and Systems, vol. 13, pp. 971–985 (2019)

27. Burton, A. et al. Wireless, battery free subdermally implantable photometry systems for chronic recording of neural dynamics. Proceedings of the National Academy of Sciences, vol. 117, pp. 2835–2845 (2020)

28. Charthad, J. Weber, M. J. Chang, T. C and Arbabian, A., A mm-Sized Implantable Medical Device (IMD) With Ultrasonic Power Transfer and a Hybrid Bi-Directional Data Link IEEE Journal of Solid-State Circuits, vol. 50, pp. 1741–1753 (2015)

29. Seo, D. et al. Wireless Recording in the Peripheral Nervous System with Ultrasonic Neural Dust Neuron, vol. 91, pp. 529–539 (2016)

30. Laiwalla, F. et al. A Distributed Wireless Network of Implantable Sub-mm Cortical Microstimulators for Brain-Computer Interfaces 2019 41^st^ Annual International Conference of the IEEE Engineering in Medicine and Biology Society (EMBC), pp 6876–6879 (2019)

31. Piech, D. K. et al. A wireless millimetre-scale implantable neural stimulator with ultrasonically powered bidirectional communication. Nature Biomedical Engineering, vol. 4, pp. 207–222, (2020)

32. Lee, S. et al. A 250 μm × 57 μm Microscale Opto-electronically Transduced Electrodes (MOTEs) for Neural Recording. IEEE Transactions on Biomedical Circuits and Systems, vol. 12, pp. 1256–1266, Dec. 2018.

33. Agrawal, D. et al. Conformal Phased Surfaces for Wireless Powering of Bioelectronic Microdevices. Nature Biomedical Engineering, vol. 1 (2017)

34. Lim, J. et al. A 0.19 × 0.17 mm2 Wireless Neural Recording IC for Motor Prediction with Near-Infrared-Based Power and Data Telemetry 2020 IEEE International Solid- State Circuits Conference - (ISSCC), pp. 416–418 (2020)

35. Singer, A. et al. Magnetoelectric Materials for Miniature, Wireless Neural Stimulation at Therapeutic Frequencies. Neuron 107, 631–643. (2020)

36. Yu, Z. et al. MagNI: A Magnetoelectrically Powered and Controlled Wireless Neurostimulating Implant. IEEE Transactions on Biomedical Circuits and Systems, 14(6), 1241–1252. (2020)

37. Nan, C. W., Bichurin, M. I., Dong, S., Viehland, D., & Srinivasan, G. Multiferroic magnetoelectric composites: Historical perspective, status, and future directions. Journal of Applied Physics, 103(3). (2008).

38. Yu, Z et al. An 8.2mm3 Implantable Neurostimulator with Magnetoelectric Power and Data Transfer 2018 IEEE International Solid - State Circuits Conference - (ISSCC) (2020)

39. Yu Z et al. Multisite bio-stimulating implants magnetoelectrically powered and individually programmed by a single transmitter 2021 IEEE Custom Integrated Circuits Conference (CICC) (2021)

40. Singer, A., & Robinson, J. T. Wireless Power Delivery Techniques for Miniature Implantable Bioelectronics. Advanced Healthcare Materials, 2100664, (2021).

41. Fang, Z. et al. Enhancing the magnetoelectric response of Metglas/polyvinylidene fluoride laminates by exploiting the flux concentration effect. Applied Physics Letters, vol. 95, p. 112903 (2009)

42. Moman, R. N. et al. Infectious Complications of Dorsal Root Ganglion Stimulation: A Systematic Review and Pooled Analysis of Incidence. Neuromodulation: Technology at the Neural Interface (2021)

43. Shen, K., & Maharbiz, M. M. Ceramic packaging in neural implants. Journal of Neural Engineering, 18(2). (2021)

44. Qanadli, S., et al. Subacute and Chronic Benign Superior Vena Cava Obstructions: Endovascular Treatment with Self-Expanding Metallic Stents. American Journal of Roentgenology 159–164. (1999)

45. Edelman, E. R., & Rogers, C. Pathobiologic responses to stenting. American Journal of Cardiology, 81(7 A), 4E–6E. (1998)

46. Stoney, W. S. et al. The Incidence of Venous Thrombosis Following Long-Term Transvenous Pacing. Annals of Thoracic Surgery, 22(2), 166–170. (1976).

47. Sabu, C., Henna, T. K., Raphey, V. R., Nivitha, K. P., & Pramod, K. Advanced biosensors for glucose and insulin. Biosensors and Bioelectronics, 141, 111201 (2019)

48. Bhave, G., Chen, J. C., Singer, A., Sharma, A., & Robinson, J. T. Distributed sensor and actuator networks for closed-loop bioelectronic medicine. Materials Today, 46, 125–135. (2021).

49. Wang, B. et al. High Bandwidth Power Electronics and Magnetic Nanoparticles for Multichannel Magnetogenetic Neurostimulation. bioRxiv (2021)

50. Russell HK. A modification of Movat’s pentachrome stain. Arch Pathol 94(2):187–191. (1972)

51. Sheehan D, Hrapchak, B. Theory and practice of histotechnology. 2nd ed. Battelle Press; (1980)

